# Genome-wide CRISPR activation screen identifies novel receptors for SARS-CoV-2 entry

**DOI:** 10.1101/2021.04.08.438924

**Authors:** Shiyou Zhu, Ying Liu, Zhuo Zhou, Zhiying Zhang, Xia Xiao, Zhiheng Liu, Ang Chen, Xiaojing Dong, Feng Tian, Shihua Chen, Yiyuan Xu, Chunhui Wang, Qiheng Li, Xuran Niu, Qian Pan, Shuo Du, Junyu Xiao, Jianwei Wang, Wensheng Wei

## Abstract

The ongoing pandemic of coronavirus disease 2019 (COVID-19) caused by severe acute respiratory syndrome coronavirus 2 (SARS-CoV-2) has been endangering worldwide public health and economy. SARS-CoV-2 infects a variety of tissues where the known receptor ACE2 is low or almost absent, suggesting the existence of alternative pathways for virus entry. Here, we performed a genome-wide barcoded-CRISPRa screen to identify novel host factors that enable SARS-CoV-2 infection. In addition to known host proteins, i.e. ACE2, TMPRSS2 and NRP1, we identified multiple host components, among which LDLRAD3, TMEM30A and CLEC4G were confirmed as functional receptors for SARS-CoV-2. All these membrane proteins bind directly to spike’s N-terminal domain (NTD). Their essential and physiological roles have all been confirmed in either neuron or liver cells. In particular, LDLRAD3 and CLEC4G mediate SARS-CoV-2 entry and infection in a fashion independent of ACE2. The identification of the novel receptors and entry mechanisms could advance our understanding of the multiorgan tropism of SARS-CoV-2, and may shed light on the development of the therapeutic countermeasures against COVID-19.

---

The outbreak of coronavirus disease 2019 (COVID-19) has caused a global health crisis. The etiologic agent of COVID-19 is acute respiratory syndrome coronavirus 2 (SARS-CoV-2), a positive-stranded betacoronavirus (*1, 2*). SARS-CoV-2 is the seventh coronavirus known to infect humans, and is the third coronavirus, after severe acute respiratory syndrome (SARS)-CoV and Middle East respiratory syndrome (MERS)-CoV, that has caused outbreaks with significant fatality rates (*3*). SARS-CoV-2 mainly infects the respiratory system, causing symptoms at the onset of disease as fever, cough, fatigue, and myalgia (*4, 5*). Moreover, COVID-19 is associated with high rates of multiorgan symptoms, such as neurological (*6*), renal (*7*), gastrointestinal (*8*), and cardiovascular (*9*) complications, indicating the broad organotropism of SARS-CoV-2.

Like SARS-CoV, SARS-CoV-2 engages human angiotensin-converting enzyme 2 (ACE2) as the receptor to enter host cells (*10*). The interaction between SARS-CoV-2 and ACE2 is mediated by the receptor-binding domain (RBD) of the SARS-CoV-2 spike (S) glycoprotein. Following binding to ACE2, S protein is cleaved into S1 and S2 domains by cellular proteases such as furin, followed by further cleavage of S2 by proteases such as TMPRSS2 or cathepsins (*11, 12*). This “priming” process triggers dramatic conformational changes of the S2 domain to enable the fusion of the viral envelope with cellular membranes, thereby allowing the release of the viral genome into host cells (*11, 13*). Despite data showing that ACE2 is a high-affinity receptor for SARS-CoV-2 (*14*), lines of evidence suggested that alternative receptors or pathways may exist. First, the tissue distribution pattern of ACE2 does not fully correlate with SARS-CoV-2 tropism, questioning the ACE2-dependent pathway as the sole entry route. Analyses of the single-cell RNA sequencing data indicated that ACE2 is expressed low throughout the respiratory tract, the primary infection site of SARS-CoV-2 (*15, 16*). Moreover, SARS-CoV-2 infects the brain, and viral RNA has been detected in immune cells such as neutrophils, macrophages, T/B cells, and NK cells (*17, 18*), whereas ACE2 is barely detected in these tissues or cells (fig. S1). Second, a recent report showed that an ACE2-null lung adenocarcinoma cell is highly permissive to SARS-CoV-2 (*19*), indicating that SARS-CoV-2 could leverage an alternative receptor for its entry. Third, a cell surface protein, AXL, has recently been reported to facilitate SARS-CoV-2 entry independently of ACE2 (*20*). NRP1 was found to function as an ACE2-dependent host factor, which is highly expressed in human pulmonary and olfactory neuronal cells of the epithelium, and could bind to S1 CendR motif of the viral spike protein (*21, 22*). Altogether, it is plausible to postulate that SARS-CoV-2 may gain its entry to host cells via alternative receptor(s) other than ACE2.

Functional genomics approaches such as CRISPR knockout screens have been conducted to search for critical host factors involved in SARS-CoV-2 infection (*23-26*). However, none of these screens could pinpoint novel receptors beyond ACE2, possibly due to the fact that such loss-of-function screens were performed based on cell types that the expression and function of ACE2 are dominant. Herein, aiming to systematically interrogate host factors for SARS-CoV-2 entry, we performed a genome-wide CRISPR activation screen in HEK293T cells using the SARS-CoV-2 spike-pseudotyped virus (*27*). Such gain-of-function screen could potentially identify those proteins that confer host cell susceptibility to SARS-CoV-2.

To establish a CRISPRa screening for the identification of viral entry factors, we utilized pseudotyped virus harboring the SARS-CoV-2 spike protein and an EGFP marker that indicates viral infection. EGFP signal was barely detectable two days after infection with different amounts of SARS-CoV-2 pseudovirus in HEK293T cells, unlike infection by lentivirus harboring the vesicular stomatitis virus G protein (VSV-G) (fig. S2A), indicating that SARS-CoV-2 pseudovirus hardly infects HEK293T cells. This was likely due to the lack of sufficient expression of functional receptors in HEK293T cells. Indeed, HEK293 cells stably overexpressing ACE2 were highly susceptible to SARS-CoV-2 pseudovirus, and the EGFP signal was proportionally boosted with the increase of pseudovirus (fig. S2B). We then tested the effect of gene activation through CRISPRa using the 50-fold concentrated pseudovirus. In HEK293T cells stably expressing CRISPRa system (HEK293T-CRISPRa cells), the upregulation of *ACE2* by sgRNA1^ACE2^ and sgRNA5^ACE2^ enabled the infection of SARS-CoV-2 pseudovirus with a significant boost of EGFP expression within the cells (Fig. 1A). As such, we developed a CRISPRa screen method to identify host factors enabling SARS-CoV-2 infection.

**Fig. 1.**
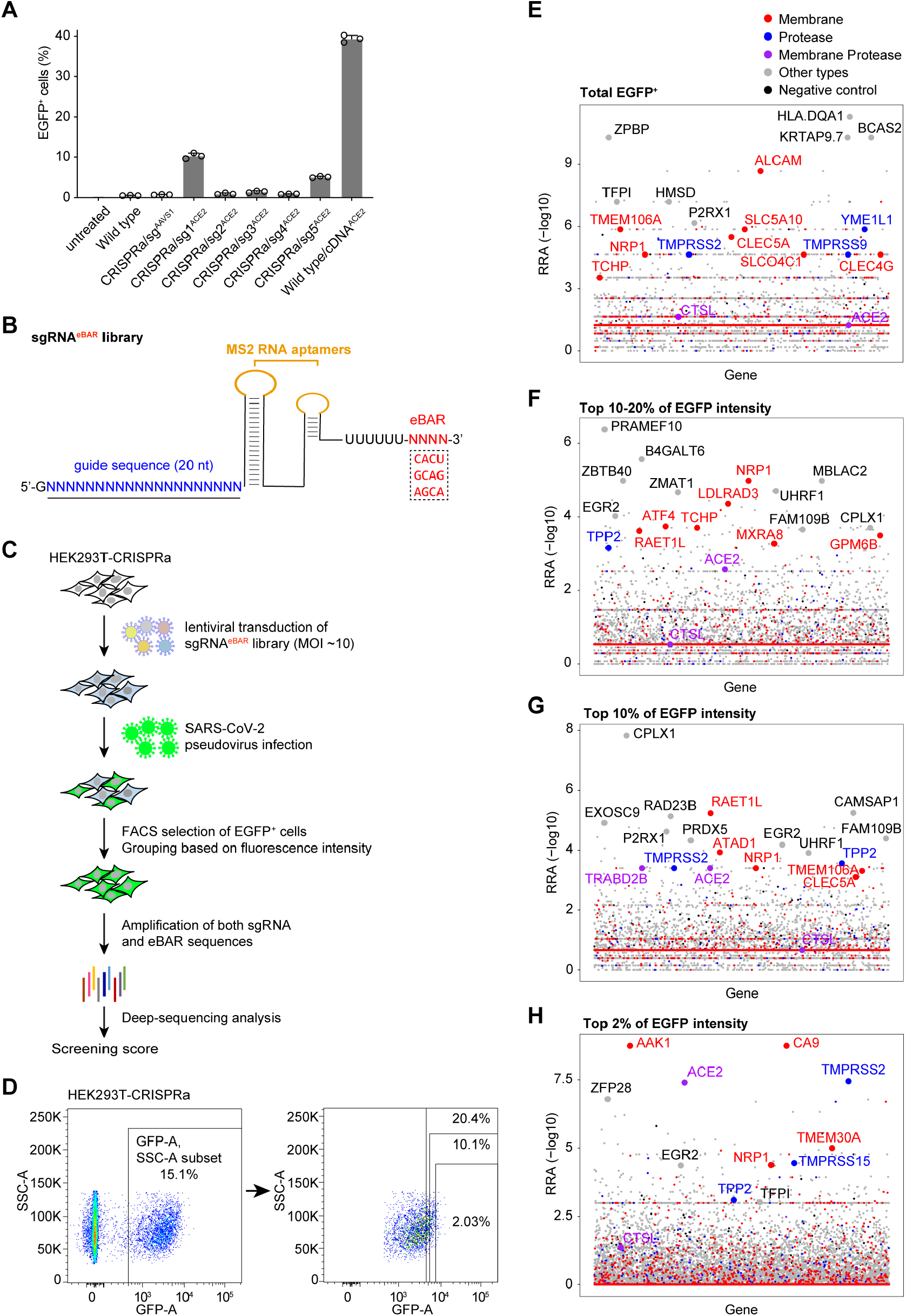
Identification of candidate factors for SARS-CoV-2 entry by a genome-wide CRISPRa gain-of-function screen in HEK293T cells. (**A**) Detection of the pseudovirus infection in HEK293T-CRISPRa cells transfected with different sgRNAs targeting *ACE2*. The infection rate of SARS-CoV-2 pseudovirus is indicated by the percentage of EGFP positive cells. Wide type/cDNA^ACE2^ represents the wild type HEK293T cells transfected with ACE2 cDNA as a positive control. (**B**) Schematic diagram of an sgRNA with an external barcode (eBAR). Three 4-nt eBARs were respectively embedded outside of the sgRNA scaffold after the poly-U signal. (**C**) Schematic of the CRISPRa screen in HEK293T cells using the SARS-CoV-2 pseudotyped virus. (**D**) FACS selection of EGFP^+^ cells grouping based on different fluorescence intensities after SARS-CoV-2 pseudovirus infection. Left indicates the total EGFP intensity of HEK293T-CRISPRa library cells after second round of pseudovirus infection. Right indicates three additional sorting gates including top 10-20%, top 10% and top 2% of the total EGFP^+^ cells. (**E**-**H**) Robust rank aggregation (RRA) scores of all genes from the total EGFP^+^ (**E**), top 10-20% (**F**), top 10% (**G**) and top 2% (H) of the total EGFP^+^ cells. RRA scores were used to evaluate the enrichment of candidate genes, which were calculated by binomial *p*-values of sgRNAs^eBAR^ targeting each gene. Membrane proteins were labelled as red dots, proteases were labelled as blue dots, the genes that are both membrane protein and proteases were labelled as purple dots. Grey and black dots represent other types of genes and negative controls.

To reach the optimal performance using the CRISPR activation system (*28*), we tended to construct a genome-wide CRISPRa library with all sgRNAs barcoded so that we could benefit from a high multiplicity of infection (MOI) in generating the cell library, an approach we previously established (*29*). Because the loops of sgRNAs are used for such a CRISPRa system we employed (*28*), we decided to add the barcodes at the external region outside of sgRNA at its 3’ end, designated as eBAR, instead of iBAR we designed before (*29*). Three external barcodes of 4-nt were assigned to each sgRNA (Fig. 1B). The oligos of sgRNA library (*30*) were synthesized and respectively cloned into three lentiviral sgRNA^eBAR^ backbones (table S1). The sgRNA^eBAR^ library was delivered into HEK293T-CRISPRa cells by lentiviral infection at an MOI of ∼10. The pseudovirus (50-fold) was added to the library cells, and the infected cells were sorted by FACS (fig. S3A). Since the EGFP signal was maintained in the sorted cells and could not completely fade out, we were not able to perform multiple rounds of enrichment to reduce noises (Fig. 1E-H). We therefore categorized screening results based on fluorescence intensity, and selected those top candidates from each group to maximize the chance of target identification. After two rounds of pseudovirus infection and sorting, we collected total EGFP^+^ cells as well as top 10-20%, top 10% and top 2% of sorted cells grouped by the EGFP intensity (Fig. 1C-D and fig. S3B). We generated screen scores for genes in each EGFP^+^ group considering the performance of all their targeting sgRNAs^eBAR^ (Fig. 1E-H, table S2, see Methods). In most groups, the known SARS-CoV-2 receptor ACE2 (*11, 13, 31*) and the main host protease TMPRSS2 (*11*) were significantly enriched. We also identified other reported host factors for SARS-CoV-2 entry, such as NRP1 (*21, 22*). The EGFP intensity was supposed to represent the strength of the target host factor in promoting virus entry. Thus we assumed that receptors were more likely to be identified from groups with higher EGFP intensity. For example, ACE2 was ranked higher in the top 2% than in other groups (Fig. 1E-H).

To further characterize these identified host factors, we performed Gene Ontology (GO) enrichment analysis (*32*). A number of genes were enriched in multiple important cellular processes, such as regulation of plasma membrane-bound cell projection organization, vesicle-mediated transport, receptor-mediated endocytosis, and viral life cycle (Fig. 2A, fig. S4, table S3). Many of these genes were top-ranked in most groups of the sorted EGFP^+^ cells (Fig. 2B). Assuming that the intensity of EGFP represented the strength of candidate factors in facilitating virus entry, we were particularly interested in membrane proteins identified from the top 2% and 10% groups. For other types of candidates, we pooled top-ranked candidates in all four groups for validation. For each gene, we found that most of its corresponding sgRNAs^eBAR^ were significantly enriched, indicating the reliability of our selection on the top hits. Besides, most of the functional sgRNAs performed consistently with their eBARs (fig. S5). The gene expression analysis revealed that several genes are widely expressed in multiple tissues such as *TMEM30A* and *CTSL*, and some genes’ expressions are more tissue-specific, such as brain-specific genes *CPLX1, LDLRAD3, GPM6B* and *EPHB1*, liver-specific genes *CLEC4G* and *MASP1*, lung-specific genes *CLEC5A* and *HLA-DQA1*, and immune-specific genes *ICAM2* and *STAMBPL1* (Fig. 2C). These findings hold the potential to interpret the organotropism of SARS-CoV-2 especially where the known receptors and other entry factors were lowly expressed.

**Fig. 2.**
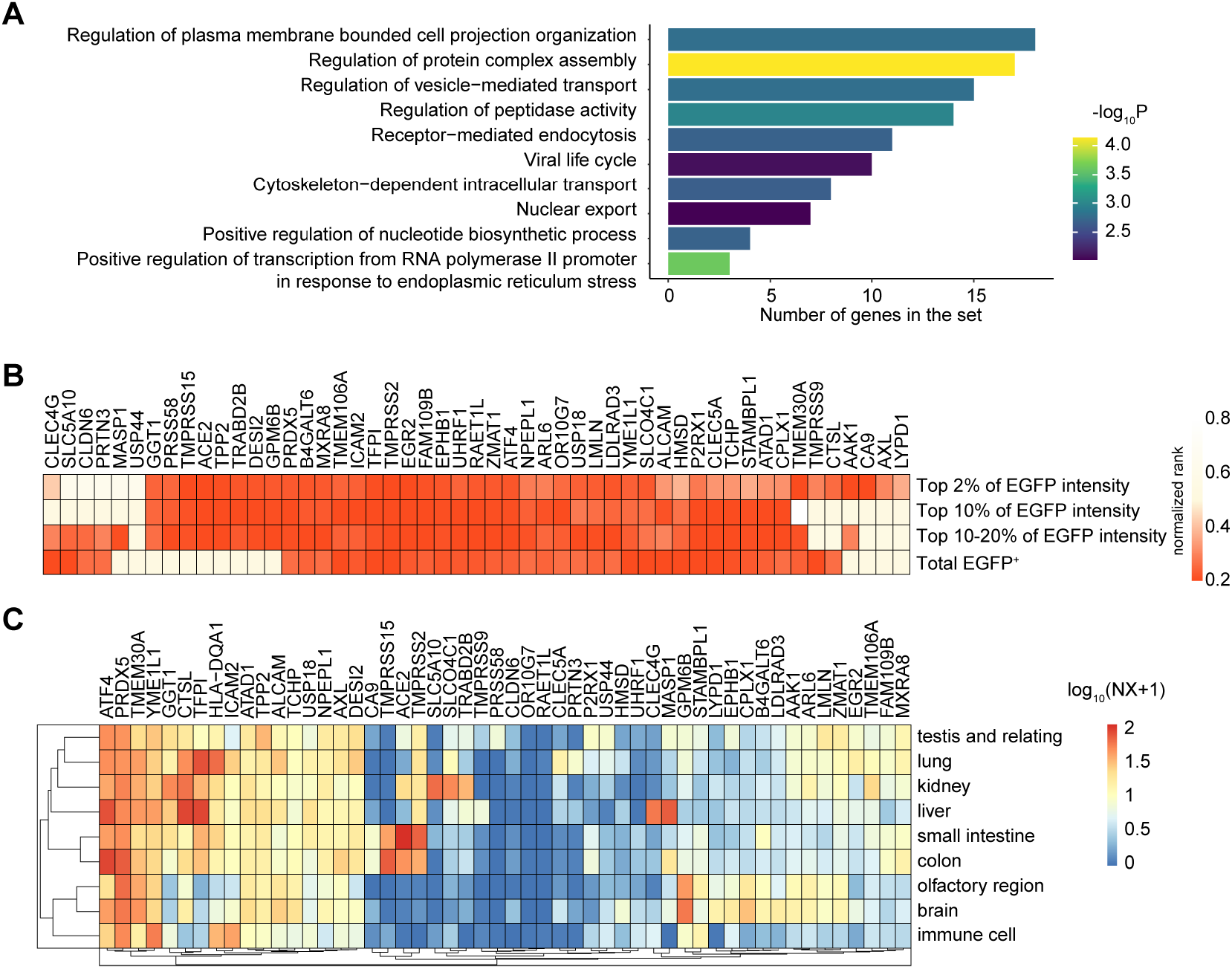
Host factors identified from CRISPRa library screening. (**A**) Gene Ontology (GO) enrichment analysis was conducted using all the significant candidates (RRA score < 0.001) identified in the four groups. Hypergeometric test was used to calculate all the *p*-values. The top-enriched GO terms were selected for visualization. The x axis represents the number of genes identified in the specific GO terms. A complete list of genes in each GO term is in table S3. (**B**) The performance of all the significant hits in four screening groups (top 2%, top 10%, top 10-20% of EGFP intensity and total EGFP^+^). The gene with a smaller value of normalized rank (in redder colour) represented a higher enrichment in the relevant groups. (**C**) Expression patterns of identified candidates within human tissues. The data used for analysis were retrieved from the Human Protein Atlas normalized expression.

To validate the candidate genes identified from our screen, we focused particularly on membrane proteins, proteases, and some other top-ranked hits. For a total of 51 candidates, we transduced HEK293T cells with their corresponding cDNAs, followed by infection with SARS-CoV-2 pseudotyped virus containing a luciferase reporter (*27*). As the known receptor or co-receptor for SARS-CoV-2, the ectopic expression of ACE2 or NRP1 greatly promoted the pseudotyped virus infection (Fig. 3A). A number of novel host factors have been confirmed to facilitate SARS-CoV-2 pseudovirus entry, including some membrane proteins, LDLRAD3, TMEM30A, CLEC4G, CPLX1, and CA9 (Fig. 3A). LDLRAD3 is a member of the LDL scavenger-receptor family that is highly expressed in neurons and has been reported to regulate amyloid precursor protein in neurons (*33*). TMEM30A is a transmembrane protein involved in membrane trafficking and signaling pathways as a heterocomplex with ATP8A1 by regulating the translocation of phospholipids (*34*). CLEC4G is a member of the C-type lectin family that has been reported to enhance the infection of SARS-CoV by interacting with its spike protein (*35*). CPLX1 is a member of the complexin/synaphin family involved in synaptic vesicle exocytosis and transmitter release (*36*). CA9, a transmembrane protein and a tumour marker (*37*), has also been reported to be involved in HBV infection (*38*). Interestingly, two proteases, STAMBPL1 and TMPRSS15, were also identified with their confirmed roles to promote SARS-CoV-2 pseudovirus infection (Fig. 3A). Proteases such as TMPRSS2 are known to play critical roles in ACE2-dependent virus entry (*11*). Therefore, we reasoned that proteases with similar functions could promote virus entry upon overexpression. We went on to validate these candidate genes using the authentic SARS-CoV-2 virus. In HEK293T cells, besides *ACE2* and *CTSL*, the ectopic expression of any of the following genes could effectively enable SARS-CoV-2 infection, *CLEC4G, CPLX1, LDLRAD3, TMEM30A*, and *STAMBPL1* (Fig. 3B).

**Fig. 3.**
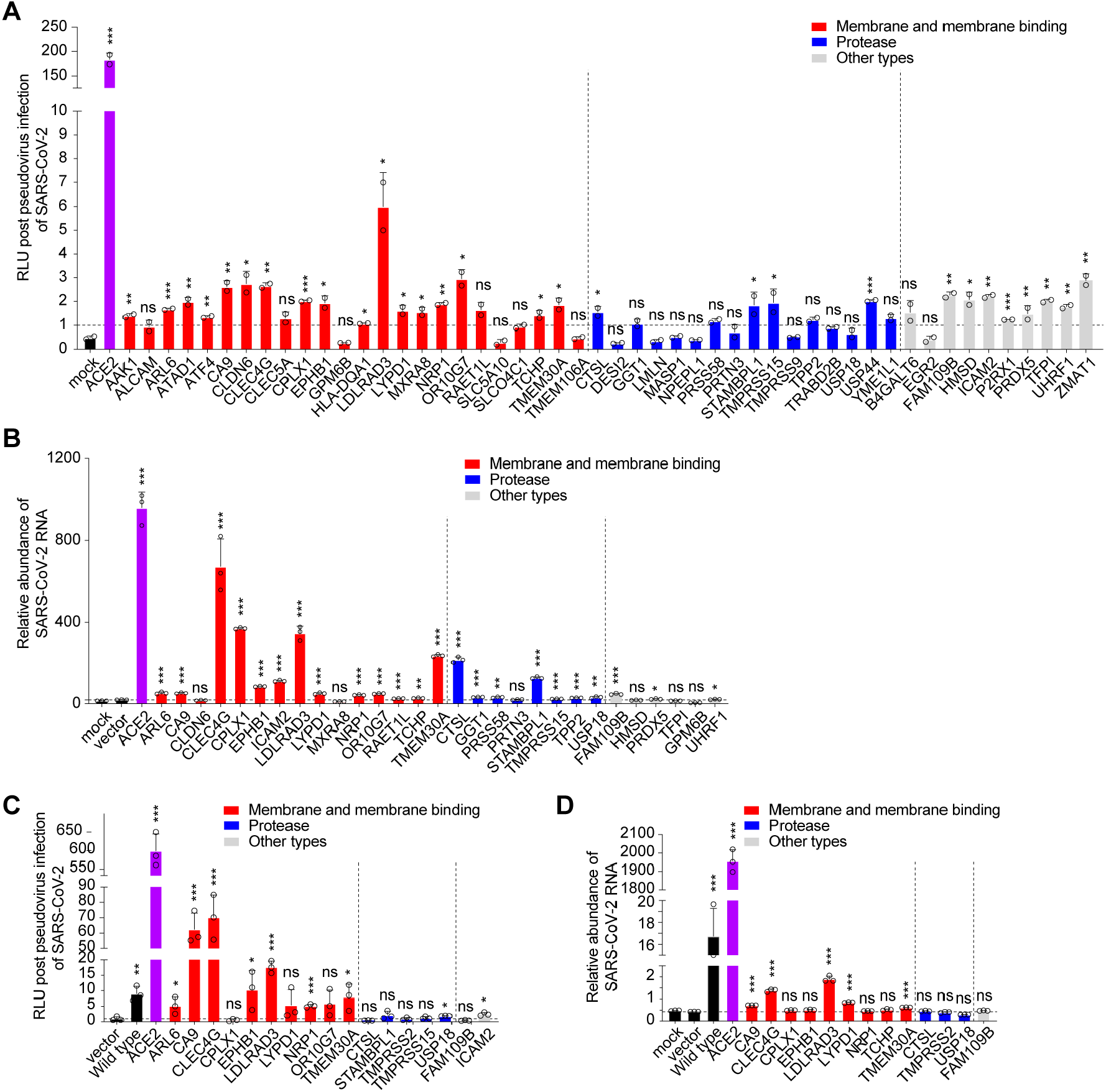
Validation of candidate genes identified from CRISPRa library screening. (**A** and **B**) Effects of identified genes on the infection of SARS-CoV-2. (**A**) 51 individual cDNAs and an empty vector were transfected into HEK293T cells. Then the cells were treated with luciferase-labelled SARS-CoV-2 pseudotyped virus. The entry of SARS-CoV-2 pseudotyped virus was quantified through measuring luciferase activity 48 h later. The luciferase activities were normalized by the empty vector. Data are presented as the mean ± s.d. (n = 2). (**B**) The cDNAs of candidate genes were introduced into HEK293T cells lentivirally labelled with an mCherry marker. The mCherry-positive cells were enriched through FACS followed by infection with authentic SARS-CoV-2 virus at an MOI of 0.5. SARS-CoV-2 RNAs were quantified by real-time qPCR and normalized by *GAPDH*. Data were presented as the mean ± s.d. (n = 3). (**C** and **D**) Effects of identified genes on the infection of SARS-CoV-2 in *ACE2*^−/–^ cells. (**C**) The cDNAs were transfected into HEK293T *ACE2*^−/–^ cells. Then the cells were treated with 10-fold concentrated SARS-CoV-2 pseudotyped virus. The entry of pseudotyped virus was quantified through measuring luciferase activity and was normalized by the empty vector. (**D**) The cDNAs of candidate genes were introduced into HEK293T *ACE2*^−/–^ cells lentivirally. Cells were enriched through FACS followed by infection with authentic SARS-CoV-2 virus at an MOI of 0.5. SARS-CoV-2 RNAs were quantified by real-time qPCR and normalized by *GAPDH*. Data were presented as the mean ± s.d. (n = 3). *P* values were calculated using Student’s *t* test, \**P* < 0.05; \*\**P* < 0.01; \*\*\**P* < 0.001; ns, not significant.

To examine if ACE2 is required for any of these candidate components to promote viral infection, we generated HEK293T *ACE2*^−/–^ cells (fig. S6). We found that the function of CA9, CLEC4G, and LDLRAD3 in facilitating luciferase reporter pseudovirus infection is independent of ACE2 (Fig. 3C). In the test of authentic virus infection, overexpression of either CLEC4G or LDLRAD3 is sufficient to enable SARS-CoV-2 infection in HEK293T *ACE2*^−/–^ cells (Fig. 3D).

Next, we focused on characterizing these membrane proteins and evaluating whether any of them serves as a functional receptor for SARS-CoV-2. We first examined whether there are interactions between these receptor candidates and SARS-CoV-2 spike (S) protein. Co-immunoprecipitation (Co-IP) assay showed that SARS-CoV-2 S co-precipitated with multiple candidate proteins including TMEM30A, ICAM2, CA9, LDLRAD3, CLEC4G, and the known host factors, ACE2, NRP1, TMPRSS2 and CTSL, but not with STAMBPL1 and CPLX1 (Fig. 4A). We then purified these proteins (fig. S7A) to examine the direct interactions by the pull-down assay. Like ACE2, LDLRAD3 and CLEC4G efficiently pulled down SARS-CoV-2 S (Fig. 4B). Reciprocally, SARS-CoV-2 S pulled down LDLRAD3, CLEC4G and ACE2, but not CA9 (fig. S7B). Moreover, we determined domains on SARS-CoV-2 S that mediate the interactions. In consistent with previous reports (*10*), ACE2 interacted with RBD but not NTD (Fig. 4D and 4C). However, NTD but not RBD were found to directly interact with LDLRAD3, CLEC4G, and TMEM30A (Fig. 4C-D).

**Fig. 4.**
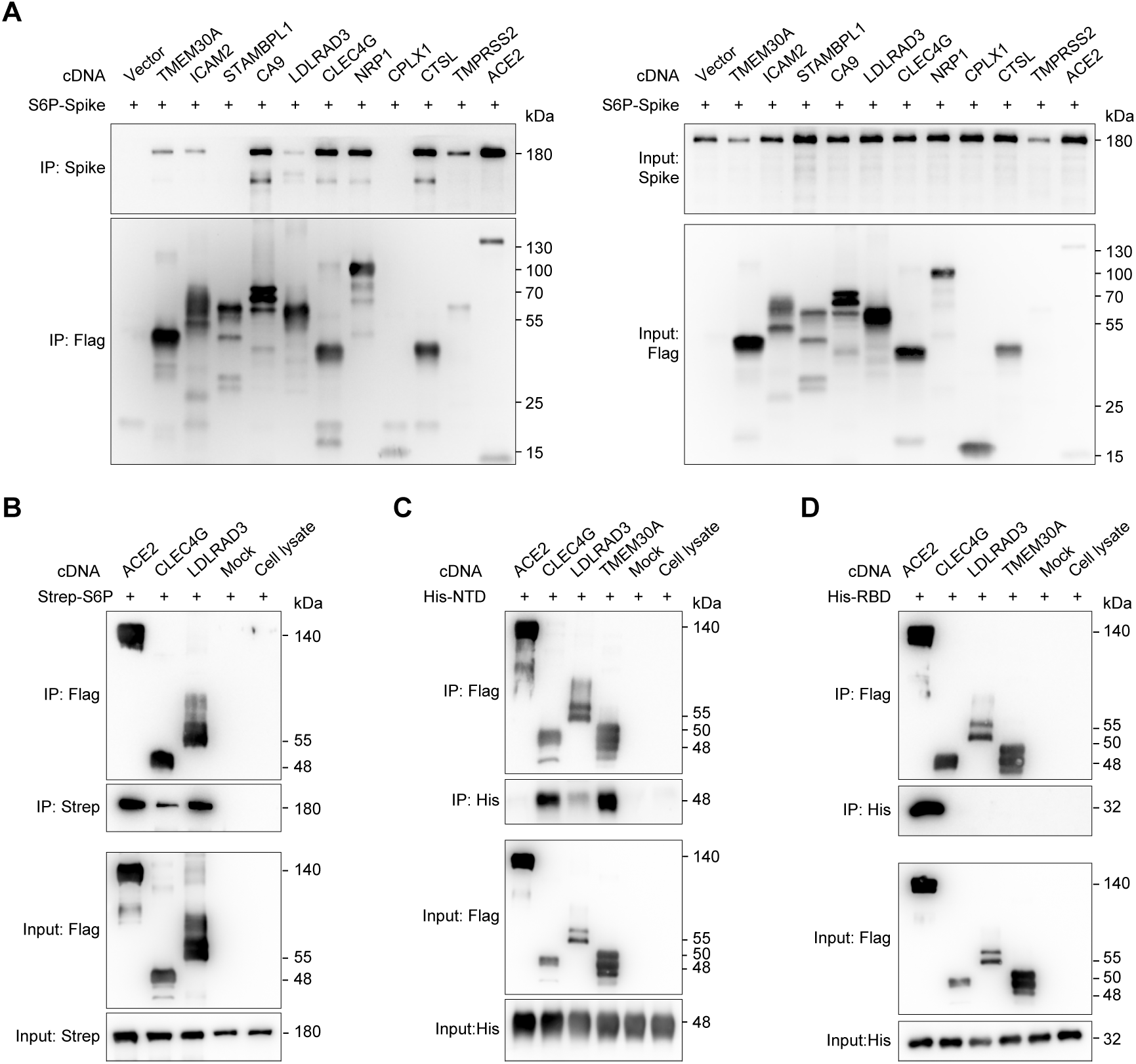
Direct binding of LDLRAD3, CLEC4G and TMEM30A to SARS-CoV-2 S. (**A**) Co-IP of SARS-CoV-2 S6P spike with FLAG-tagged proteins in HEK293T cells transfected with Flag-cDNA constructs and SARS-CoV-2 S. Immunoblot analysis was conducted using anti-Flag and anti-spike antibodies. (**B**) *In vitro* pull-down assay of purified ACE2, CLEC4G and LDLRAD3 to SARS-CoV-2 S. Strep-tagged SARS-CoV-2 S and FLAG-tagged full-length candidate receptors were expressed in HEK293T cells and affinity-purified. Immunoblot analysis was conducted using anti-Flag and anti-Strep antibodies. (**C** and **D**) *In vitro* pull-down assay of purified ACE2, CLEC4G, LDLRAD3 and TMEM30A to the NTD (**C**) or RBD (**D**) of SARS-CoV-2 S. Immunoblot analysis was conducted using anti-Flag and anti-His antibodies.

In light of these direct binding results, we predicted that the extracellular addition of these purified proteins could prevent virus entry by competing cellular receptors for binding to S. To test this idea, we incubated serially diluted soluble proteins with authentic SARS-CoV-2 virus before infection. The addition of soluble ACE2 (Fig. 5A and 5B) and LDLRAD3 (Fig. 5C and 5D) were capable of protecting both SH-SY5Y (Fig. 5A and 5C) and SK-N-SH (Fig. 5B and 5D), two neuroblastoma cell lines, from SARS-CoV-2 infection, in a dose-dependent manner. Similarly, the addition of soluble ACE2 (Fig. 5E) and CLEC4G (Fig. 5F) effectively suppressed SARS-CoV-2 infection in a liver cancer cell line Huh7.5, also in a dose-dependent manner.

**Fig. 5.**
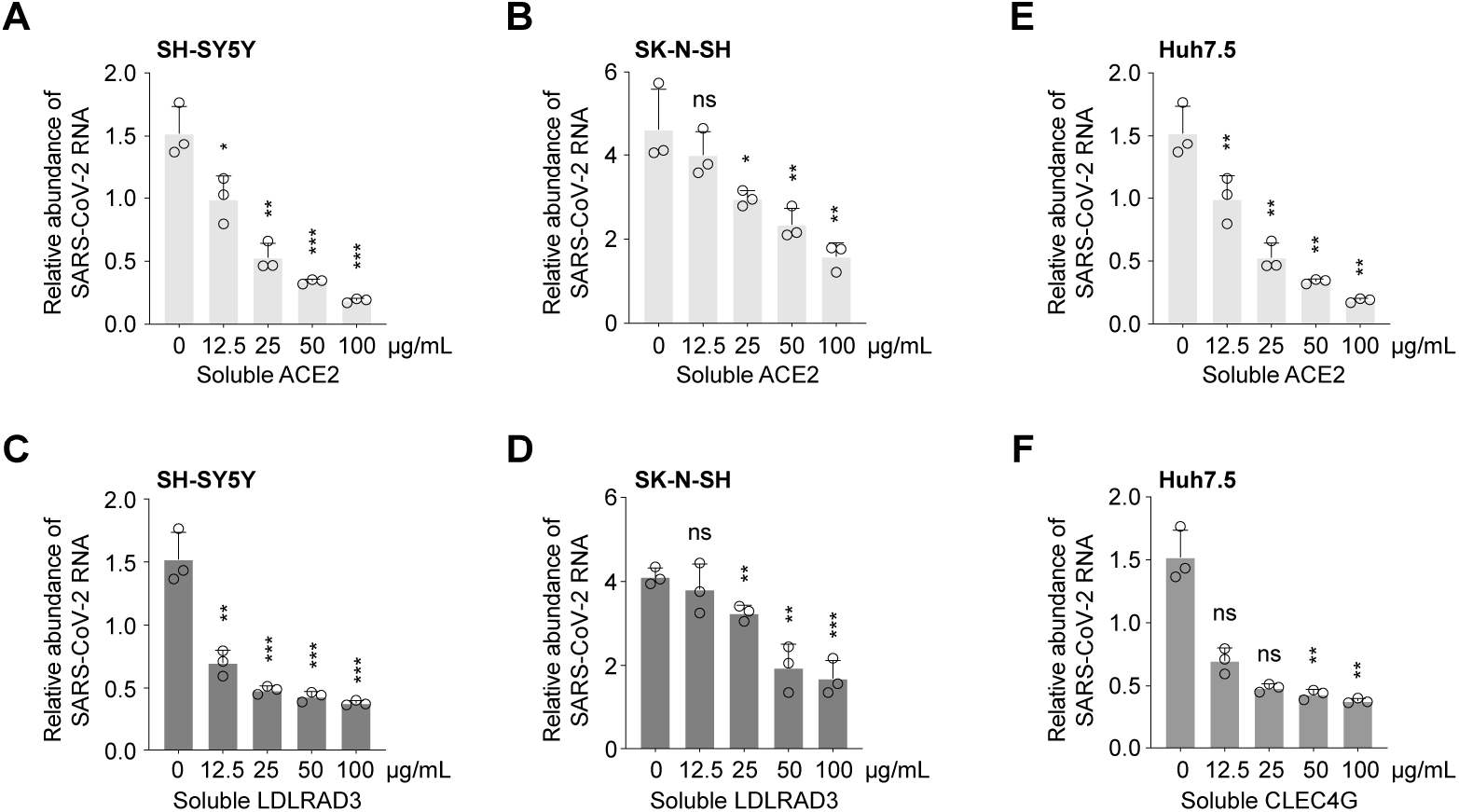
Inhibition of soluble proteins on SARS-CoV-2 infection. (**A** and **B**) Effects of purified ACE2 on SARS-CoV-2 infection in SH-SY5Y (**A**) and SK-N-SH (**B**) cells. (**C** and **D**) Effects of purified LDLRAD3 on SARS-CoV-2 infection in SH-SY5Y (**C**) and SK-N-SH (**D**) cells. (**E** and **F**) Effects of purified ACE2 (**E**) and CLEC4G (**F**) on SARS-CoV-2 infection in Huh7.5 cells. The soluble proteins (0, 12.5, 25, 50 and 100 µg/mL) were incubated with authentic SARS-CoV-2 virus for 1 h. Infection was performed at an MOI of 0.5. SARS-CoV-2 RNAs were quantified by real-time qPCR and normalized by *GAPDH*. Data were presented as the mean ± s.d. (n = 3). *P* values were calculated using Student’s *t* test, \**P* < 0.05; \*\**P* < 0.01; \*\*\**P* < 0.001; ns, not significant.

SARS-CoV-2’s entry is initiated by the interaction between the spike and its host receptor(s), followed by furin-mediated cleavage at the S1/S2 site and the priming via TMPRSS2 or other surface/endosomal proteases (*39, 40*). The surface subunit S1 of spike is responsible for binding to the host receptor, and the transmembrane subunit S2 mediates the viral and cellular membrane fusion (*11*). Previous studies have shown that SARS-CoV-2 S present in the plasma membrane possesses high fusogenic activity and could trigger receptor-dependent fusion with neighboring cells, leading to the formation of multinucleated giant cells (syncytia) (*40, 41*). To examine whether the interaction between the SARS-CoV-2 spike protein and our newly identified receptors could elicit membrane fusion, we performed a co-culture assay to determine the syncytium formation. The wild-type HEK293T cells transfected with plasmids expressing S and EGFP were mixed with HEK293T cells stably overexpressing individual candidate receptors labelled with an mCherry marker (see Methods). At 40 h post cell co-culture, cells expressing any of the following, *ACE2, CLEC4G, LDLRAD3* and *TMEM30A*, substantially fused with cells expressing S, manifested by the colocalization of the EGFP and mCherry fluorescent signals in the merged images, which could be visualized even in the bright field (Fig. 6). In comparison, the control cells infected with only the empty vector showed no syncytium formation, nor merged fluorescent signals (Fig. 6). These observations suggested that any of CLEC4G, LDLRAD3 and TMEM30A functionally interacts with spike protein of SARS-CoV-2 S to trigger membrane-to-membrane fusion, a critical step for receptor-mediated viral entry, just as ACE2.

**Fig. 6.**
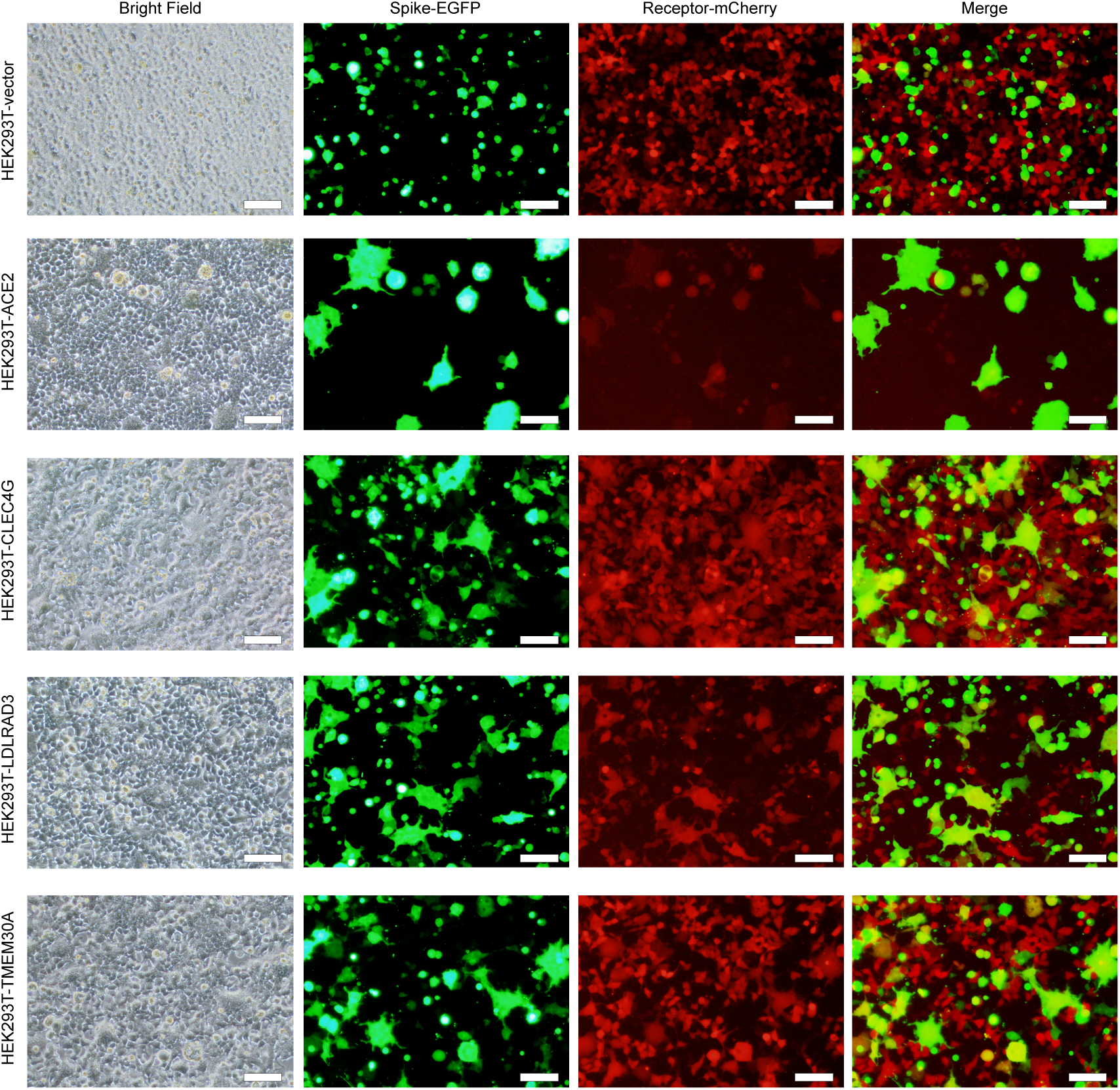
Examination of the interaction between SARS-CoV-2 S protein and candidate receptors by syncytium formation assay. Spike-EGFP represents the HEK293T cells transfected with SARS-CoV-2 S protein and an EGFP marker. Receptor-mCherry represents the HEK293T cells stably overexpressed with the known and candidate receptors labelled with an mCherry marker, labeled as HEK293T-ACE2, HEK293T-CLEC4G, HEK293T-LDLRAD3 and HEK293T-TMEM30A. HEK293T cells infected with the cDNA-expressing vector, labeled as HEK293T-vector, was served as the control. Merge indicates the co-localization of the two categories of cells through merging the EGFP and mCherry fluorescence channels by ImageJ. The images were taken 40 h after co-culturing the two categories of cells. The scale bar = 100 µm.

To evaluate their physiological roles, we first conducted expression analysis using the Human Protein Atlas (HPA) (*42*). LDLRAD3 is preferentially expressed in brain tissue, such as cerebellum, spinal cord and salivary gland (fig. S8A). The expression of CLEC4G could only be detected in liver and lymph node (fig. S8B). While TMEM30A is more ubiquitously expressed in tissues including those with a high incidence of infection, such as lung, colon and airway (fig. S8C). We then tested whether these candidate receptors are required for SARS-CoV-2 in specific cells. *TMEM30A* and *LDLRAD3* showed much higher expression compared to *ACE2* in SH-SY5Y cells, which is consistent with the analysis from HPA (Fig. 7A). The siRNAs targeting *ACE2, TMEM30A, LDLRAD3* were introduced into indicated cells followed by authentic SARS-CoV-2 infection (table S4). Efficient knockdown was confirmed by qPCR analysis (Fig. 7B-D). The disruption of *ACE2, LDLRAD3* and *TMEM30A* expression all led to significant cellular resistance to SARS-CoV-2 infection (Fig. 7E). Of note, the extent of siRNA knockdown correlated well with the inhibitory effects to the viral infection (Fig. 7E). Similar in SH-SY5Y, *LDLRAD3* and *TMEM30A* were highly expressed in another neuron cell line, SK-N-SH (Fig. 7F). Moreover, the expression of *ACE2* in SK-N-SH was too low to be detected through qPCR. The siRNA knockdown of *LDLRAD3* and *TMEM30A* (Fig. 7G and H) significantly blocked SARS-CoV-2 infection, whereas *ACE2*-targeting siRNAs exerted no effects, likely due to the lack of endogenous *ACE2* expression (Fig. 7I). As CLEC4G is preferentially expressed in the liver, we tested its function in Huh7.5 cells. The qPCR results indicated a lower expression level of *CLEC4G* than *ACE2* in Huh7.5 (Fig. 7J). Nevertheless, knockdown of either *ACE2* or *CLEC4G* (Fig. 7K and L) significantly inhibited SARS-CoV-2 infection in Huh7.5 cells (Fig. 7M). These results clearly demonstrated the essential roles of LDLRAD3, TMEM30A and CLEC4G in SARS-CoV-2 infection, especially in cell types where ACE2 was lowly expressed.

**Fig. 7.**
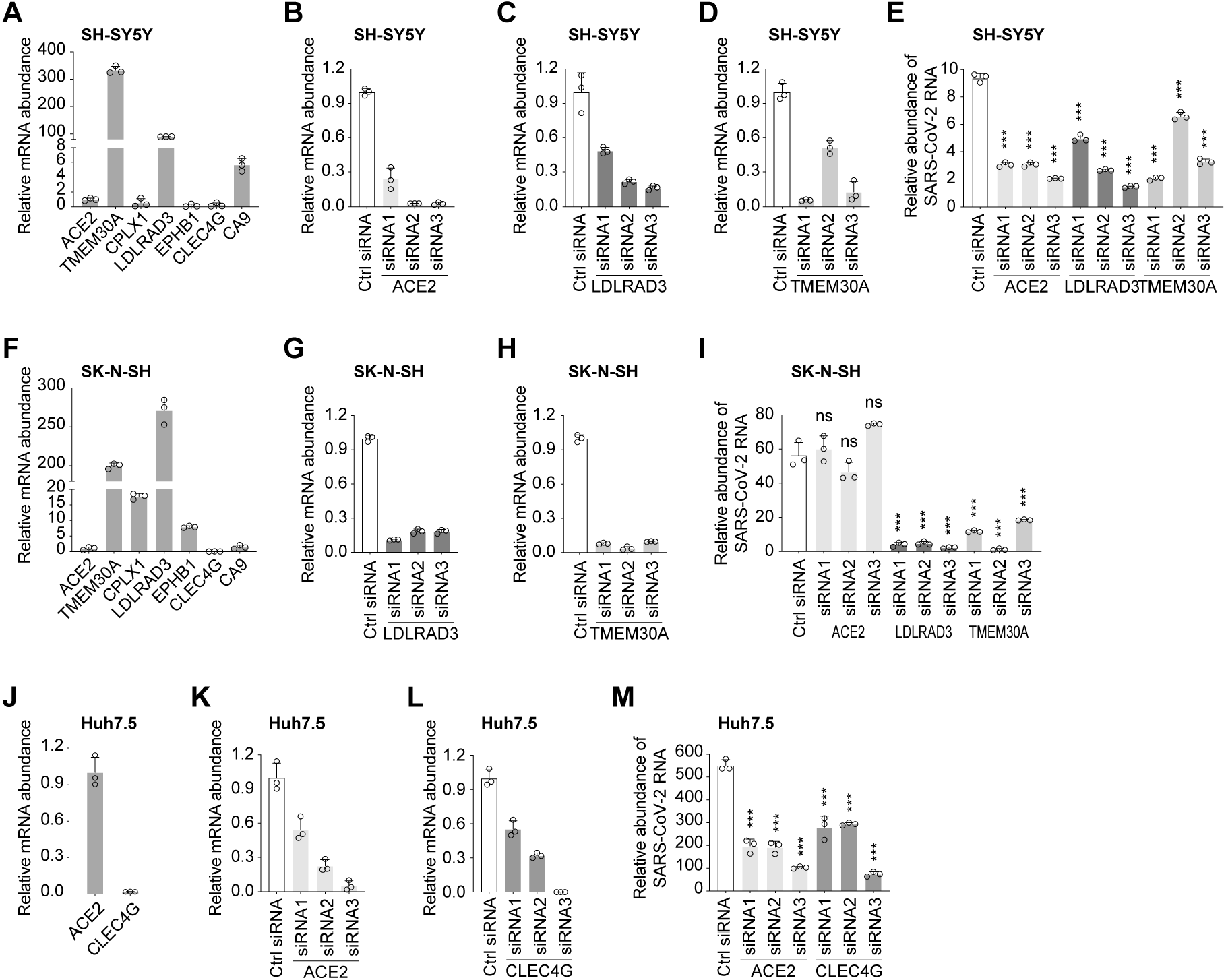
Loss-of-function effects of identified receptors on SARS-CoV-2 infection. (**A**) Expression of identified host factors relative to *ACE2* in SH-SY5Y cells. (**B** to **D**) Suppression of *ACE2* (**B**), *LDLRAD3* (**C**) and *TMEM30A* (**D**) by siRNAs in SH-SY5Y cells. (**E**) Effects of suppression of candidate receptors by siRNAs on SARS-CoV-2 infection in SH-SY5Y cells. Infection was performed at an MOI of 0.5. (**F**) Expression of identified host genes relative to *ACE2* in SK-N-SH cells. (**G** and **H**) Suppression of *LDLRAD3* (**G**) and *TMEM30A* (**H**) by siRNAs in in SK-N-SH cells. (**I**) Effects of suppression of candidate genes by siRNAs on SARS-CoV-2 infection in SK-N-SH cells. Infection was performed at an MOI of 0.5. (**J**) Expression of identified host genes relative to *ACE2* in Huh7.5 cells. (**K** and **L**) Suppression of *ACE2* (**K**) and *CLEC4G* (**L**) by siRNAs in in Huh7.5 cells. (**I**) Effects of suppression of candidate genes by siRNAs on SARS-CoV-2 infection in Huh7.5 cells. Infection was performed at an MOI of 0.5. For all these experiments, a total of 20 pmol for each siRNA was transfected into cells. The relative mRNA abundance was quantified 48 h post transfection. Ctrl RNA: Random non-targeting siRNA. RNA abundance of host factors and SARS-CoV-2 were quantified by real-time qPCR and normalized by *GAPDH*. Data were presented as the mean ± s.d. (n = 3). *P* values were calculated using Student’s *t* test, \**P* < 0.05; \*\**P* < 0.01; \*\*\**P* < 0.001; ns, not significant. Primers used for real-time qPCR were listed in table S5.

Here we conducted a study of applying a gain-of-function screen for SARS-CoV-2 entry, which uncovered three new viral receptors besides ACE2. Of the candidate receptors identified in this study, LDLRAD3 is highly expressed in neurons, and its overexpression robustly enhanced SARS-CoV-2 infection in both wild-type and HEK293T *ACE2*^−/–^ cells. LDLRAD3 has recently received attention as a critical receptor for the Venezuelan equine encephalitis virus (VEEV) (*43*). Similar to VEEV, SARS-CoV-2 was also reported to infect the brain (*7*). Our data revealed that knockdown of LDLRAD3 or supplement of its soluble protein could dramatically reduce SARS-CoV-2 infection in neuron cells, suggesting its critical function in mediating viral entry in neurons. Moreover, another confirmed ACE2-independent candidate receptor, CLEC4G, was known to be highly expressed in the liver (*44*), lymph node and monocytes (fig. S8B). This gene encodes a glycan-binding receptor and is a member of the C-type lectin family, which has been found to facilitate SARS-CoV attachment through glycan-binding (*35*). Herein, we demonstrated for the first time CLEC4G’s role in SARS-CoV-2 entry. Interestingly, the transmembrane protein TMEM30A was also identified in a recent genome-wide knockout screen for SARS-CoV-2 in Huh7.5 cells (*23*). Our study confirmed its essentiality for SARS-CoV-2 entry and its direct binding with SARS-CoV-2 S. The discovery of multiple receptors in this study, with either tissue-specific or broad-spectrum expression patterns, might provide clues for understanding the multiorgan tropism of SARS-CoV-2 (*7*).

It’s worth noting that all these receptors we identified bind to the NTD domain of S, rather than RBD, the ACE2-binding domain, suggesting that NTD of SARS-CoV-2 S also plays important roles in mediating virus entry. Recent reports showed that NTD-specific neutralization antibodies isolated from convalescent COVID-19 patients enabled robust protection from the SARS-CoV-2 challenge (*45, 46*). It is possible that NTD-targeting antibodies might function by blocking NTD mediated virus entry.

Besides membrane proteins, we have also discovered several proteases, i.e., STAMBPL1 and TMPRSS15, whose overexpression promoted SARS-CoV-2 infection. The synergy of receptors and proteases in different tissues is worth of further investigation. Finally, the novel identified receptors or other functional factors brought a more comprehensive understanding for SARS-CoV-2 infection and might serve as novel therapeutic targets for COVID-19.

## Materials and Methods

### Plasmids

The lentiviral sgRNA^eBAR^-expressing backbone was constructed by inserting sgRNA scaffold embedded MS2 loops at tetraloop and stemloop 2 along with eBAR sequence into pLenti-sgRNA-Lib (Addgene, 53121). The sgRNA-expressing sequences were cloned into the backbone using the BsmBI-mediated Golden Gate cloning strategy (*47*). The pLenti-EGFP used for pseudovirus production was constructed by cloning EGFP sequence into pLenti-SV40-mCherry. The cDNA-expressing plasmids were constructed by inserting each cDNA sequence into the multiple cloning sites before the Flag tag of the pLenti-SV40-mCherry vector following the standard cloning protocol. The plasmids lenti dCAS-VP64_Blast (Addgene, 61425) and lenti MS2-P65-HSF1_Hygro (Addgene, 61426) were purchased from Addgene. The oligos of CRISPRa library were synthesized in Synbio Technologies according to the Human Genome-wide CRISPRa-v2 Libraries (Addgene, 83978) (*30*).

### Cell culture

The HEK293T cell line was from EdiGene Inc., and Huh 7.5 cell line was from S. Cohen’s laboratory (Stanford University School of Medicine). All these cells were maintained in Dulbecco’s modified Eagle’s medium (DMEM; Gibco, C11995500BT) supplemented with 10% fetal bovine serum (FBS; Biological Industries, 04-001-1ACS) and 1% penicillin/streptomycin, and cultured with 5% CO2 at 37°C. Sf21 insect cells were maintained in SIM SF medium (Sino Biological, RZ13NO0801) and 1% penicillin/streptomycin (Gibco, 2257215) with 110 rpm at 27°C. All cells were routinely checked to confirm the absence of mycoplasma contamination.

### Production and infection of SARS-CoV-2 pseudotyped virus

HEK293T cells were seeded 24 h before pseudovirus packaging. The SARS-CoV-2 pseudotyped virus was generated by co-transfection of the pCAGGS-S with the viral packaging plasmid psPAX2 and pLenti-EGFP/luciferase-expressing plasmid as a proportion of 1:1:1 into HEK293T cells using the X-tremeGENE HP DNA transfection reagent (Roche, 06366546001) according to the manufacturer′s instructions. The cell supernatant containing pseudovirus was collected 48 h post transfection, and was directly concentrated in different ratios using Lenti-X™ Concentrator (Clontech, 631232). The concentrated pseudovirus was immediately added into cells for infection without freez-thawing. For infection with SARS-CoV-2 pseudotyped virus, cells were seeded 24 h before virus collecting. Concentrated pseudovirus was added into culturing medium with polybrene (8 µg/mL). After 24 h, the medium was changed by conventional medium and cells were incubated for another 48 h.

### Construction of the CRISPRa sgRNA^eBAR^ plasmid library

The synthesized oligo pool of CRISPRa library was PCR amplified with primers (table S5) including the BsmBI recognition sites using Phusion®High-Fidelity PCR Kit (NEB, E0553L). After purification with DNA Clean & Concentrator-25 (Zymo Research Corporation, D4034), the purified PCR product was respectively inserted into the three sgRNA^eBAR^-expressing backbones constructed above through the Golden Gate cloning strategy (*47*). The ligation mixture of each group was separately purified with DNA Clean & Concentrator-5 (Zymo Research Corporation, D4014), and was electro-transformed into E.coli HST08 Premium Electro-Cells (Takara, 9028) according to the manufacturer’s protocol using a Gene Pulser Xcell (BioRad). Transformed clones were counted to ensure at least 300-fold coverage for each sgRNA^eBAR^. The plasmid of each sgRNA^eBAR^ library was extracted using an EndoFree Plasmid Maxi Kit (QIAGEN, 12362), and further mixed in a 1:1:1 molar ratio. The library lentivirus was generated by co-transfection of the library plasmid mixture with two lentiviral packaging plasmids pR8.74 and pVSV-G (Addgene, 12259) as a proportion of 10:10:1 into HEK293T cells. The cell supernatant containing lentivirus was collected 48 h post transfection and stored at -80°C.

### CRISPRa screening for SARS-CoV-2 entry factors

The HEK293T cells were engineered to stably express the CRISPRa system including lenti dCAS-VP64_Blast and lenti MS2-P65-HSF1_Hygro vectors, termed as HEK293T-CRISPRa cells. The HEK293T-CRISPRa cells were seeded 24 h post lentiviral infection, and were further infected with the library lentivirus at an MOI of 10 with a high coverage (5000-fold) for each sgRNA. Two days post lentiviral infection, the library cells were subjected to puromycin selection for 48 h (1 µg/mL). After puromycin treatment, the library cells were collected as the reference sample and were continuously cultured for 5 days. The fresh SARS-CoV-2 pseudovirus with EGFP marker (50-fold) was added to the library cells, and the EGFP positive cells were sorted by FACS 48 h post first round of pseudovirus infection. After culturing the sorted cells for additional several days, a second round of pseudovirus infection were conducted as described above, and the library cells were sorted for total EGFP^+^ cells as well as the top 10-20%, top 10% and top 2% grouped by the EGFP intensity. The reference sample and each group of EGFP^+^ cells were subjected to genomic extraction (QIAGEN, 69506), PCR amplification of the sgRNA^eBAR^ sequences (KAPA, KK2625) and high-throughput sequencing as previously described (*48*).

### Analysis of CRISPRa screening results

In order to calculate the enriched genes after CRISPRa screening, we developed an analysis algorithm eBAR-analyzer, which was implemented using R and could be obtained from https://github.com/wolfsonliu/FluorescenceSelection. In principle, the eBAR-analyzer algorithm adopt binomial distribution, in which the selection of cells hosting sgRNAs targeting specific genes enriched by FACS was considered as results of series of Bernoulli trials. The normalization of raw counts of sgRNAs^eBAR^ was calculated based on the cell proportion of EGFP intensity groups compared with the initial cell population. For instance, when we selected the top 2% intensity of EGFP^+^ cells, the normalization factor for the group will be 0.02. Then the normalized counts would be total detected reads multiplied by group normalization factor. The final normalization process would ensure that the smallest normalized counts will be an integer after rounded. Based on this, the *p*-values of the sgRNAs^eBAR^ were calculated by assuming counts of each intensity group were drawn from the initial population counts satisfying a binomial distribution. The normalized ranks of *p*-values for each sgRNA^eBAR^ were calculated. Finally, Robust Rank Aggregation (RRA) (*49*) was used to calculate the rank in gene level from the normalized ranks of *p*-values of sgRNAs^eBAR^. The RRA scores were the output results of the algorithm.

### GO enrichment and expression pattern analysis

The Gene Ontology (GO) enrichment analysis of identified host factors (RRA score < 0.001) was performed using Metascape Resource (*32*). Hypergeometric test was used to calculate all the *p*-values for all the terms. We selected top-enriched GO terms for visualization in this manuscript. For the expression pattern analysis of identified candidates, we use data retrieved from Human Protein Atlas to obtain normalized expression of each factor (*42*).

### Validation of identified candidates

For the individual validation of screening results, we introduced cDNAs of candidate genes into cells. The cDNAs were transfected into cells using the X-tremeGENE HP DNA transfection reagent (Roche, 06366546001). Then the transfected cells were infected by concentrated SARS-CoV-2 pseudotyped virus 48 h later. The infection of pseudotyped virus was quantified through measuring luciferase activities.

The lentiviral particles expressing individual cDNA labelled with an mCherry marker were generated by co-transfection of the cDNA plasmid mixture with two lentiviral packaging plasmids pR8.74 and pVSV-G (Addgene, 12259) as a proportion of 10:10:1 into HEK293T cells, followed by infection into cells. The cDNA transduced cells were selected through FACS and were infected with authentic SARS-CoV-2 virus at an MOI of 0.5 for 1 h. Infected cells were cultured for another 24 h with conventional medium, then treated with Trizol. The infection of authentic SARS-CoV-2 virus was quantified by real-time qPCR of RNA abundance.

### Protein production and purification

The SARS-CoV-2 NTD (residues 13-303) or RBD (residues 319-541) with a C-terminal His tag was cloned into a modified pFastBac vector (Invitrogen) that encodes a melittin signal peptide before the NTD or RBD. Bacmids DNA were generated using the Bac-to-Bac system. Baculoviruses were generated and amplified using the Sf21 insect cells, and were subsequently used to infect High Five insect cells for protein expression. NTD/RBD was retrieved from the conditioned cell growth media using the Ni-NTA resin and further purified using a Superdex 200 Increase 10/300 gel filtration column in 20 mM HEPES pH 7.2, and 150 mM NaCl. Strep-tagged S6P spike protein was expressed in the HEK293F cells and purified as described previously (*50*).

### Co-IP

For the Co-IP assay, the plasmids of SARS-CoV-2 S6P spike and individual cDNA were transfected into HEK293T cells. After 48 h, the cells were washed using precooled PBS for 3 times, then lysed with precooled lysis buffer (50 mM Tris-HCl pH 7.4, 150 mM NaCl, 0.25% Na-deoxycholate, 1 mM EDTA, 1% NonidetP-40) with Protease Inhibitor Cocktail Tablet (Thermo, VJ313124) at 4 °C for 1 h before being subjected to centrifugation at 15, 000 g at 4°C for 15 min. We transferred 30 µL sample into a new tube as the input. The rest of cell lysates were treated with Anti-Flag M2 Affinity Gel (Sigma, A2220) at 4°C overnight. Then the lysates were washed using wash buffer (50 mM Tris-HCl pH 7.4, 150 mM NaCl, 0.25% Na-deoxycholate, 1 mM EDTA, 0.1% NonidetP-40) for at least 4 times. The proteins were eluted using 250 µg/mL Flag peptide into wash buffer for 1 h at 4°C and subjected to immunoblotting analysis using antibodies for Flag tag (SIGMA, SLCD6338) and SARS-CoV-2 S (Sino Biological, 40589-T62).

### Flag pull-down assay

Potential receptor proteins with C-terminal Flag tag were transiently expressed in HEK293F cells using polyethylenimine (PEI, Polysciences). 36 h following transfection, the cells were collected by centrifugation and disrupted on ice using a dounce homogenizer in the lysis buffer [25 mM Tris pH 8.0, 150 mM NaCl, 1.0% (w/v) N-dodecyl β-d-maltoside (DDM), and 0.1% (w/v) cholesterol hemisuccinate (CHS)], supplemented with Protease Inhibitor cocktail (Bimake, B14001). After ultracentrifugation (45,000 g, 30 min, 4°C), the supernatants were first incubated with the anti-Flag affinity beads (Smart-Lifesciences, SA042025) for 2 h at 4°C with rotation. The beads were then pelleted and washed for five times with the wash buffer [25 mM Tris pH 8.0, 150 mM NaCl, 0.3% (w/v) DDM, and 0.03% (w/v) CHS]. Afterwards, the beads were incubated with purified S6P/NTD/RBD proteins as described above for 1 h with rotation. The beads were then again pelleted and washed five times with the wash buffer. Bound proteins were eluted from the beads using the elute buffer [25 mM Tris pH 8.0, 150 mM NaCl, 0.1% (w/v) DDM, 0.01% (w/v) CHS, and 250 ng/mL Flag peptide], and analyzed by immunoblotting using antibodies for the Strep tag (HuaxingBio, HX1816) or His tag (TransGen, HT501).

### StrepTactin pull-down assay

For the StrepTactin pull-down assay, potential receptor proteins were purified using the anti-Flag affinity beads and eluted as described above. Then they were incubated with purified S6P on ice for 1 h. The mixtures were then incubated with the StrepTactin beads (Smart Lifesciences) in the wash buffer at 4°C for another 1 h with rotation, washed by five times with the wash buffer, and eluted using the final buffer [25 mM Tris pH 8.0, 150 mM NaCl, 0.1% (w/v) DDM, 0.01% (w/v) CHS, and 10 mM desthiobiotin]. The results were analyzed by immunoblotting using antibodies for the Flag tag (SIGMA, SLCD6338).

### Syncytium formation assay

HEK293T cells were first transfected with the pCAGGS-S plasmid with an EGFP selection marker. 24 h post transfection, the transfected cells were detached, and mixed with HEK293T cells stably overexpressing different cDNAs (ACE2, CLEC4G, LDLRAD3, TMEM30A and the cDNA-expressing empty vector as the negative control) labelled with an mCherry marker in a 1:1 ratio. Then the two categories of cells were co-cultured in 12-well plates at about 60% confluency. After 40 h of cell co-culture, the images were captured by fluorescence microscopy.

### Real-time qPCR

The cultured cells transfected with siRNAs or/and infected with authentic SARS-CoV-2 virus were treated by Trizol. RNA was extracted using Direct-zol RNA kit (Zymo, R2069), and the cDNA was synthesized using QuantScript RT kit (TIANGEN, KR103-03). Real-time PCR was performed using SYBR Premix Ex Taq II (TaKaRa, RR820A) on LightCycler96 qPCR system (Roche). The relative RNA abundance of candidate factors or SARS-CoV-2 virus was measured and normalized by GAPDH. All the primers used for real-time qPCR were listed in table S5.

### Inhibition of SARS-CoV-2 infection by soluble proteins and siRNAs

For inhibition by soluble proteins, the purified protein of ACE2, LDLRAD3 or CLEC4G with different doses (0, 12.5 µg/mL, 25 µg/mL, 50 µg/mL and 100 µg/mL) were incubated with authentic SARS-CoV-2 virus for 1 h followed by infection at an MOI of 0.5. For inhibition by siRNAs, cells were seeded at 24-well plates 24 h before transfection. Each siRNA including negative control siRNA at an amount of 20 pmol was transfected into cells with 6 µL Lipofectamine RNAiMAX Reagent (Life technologies, 13778-150). 24 h later, the cells were infected with authentic SARS-CoV-2 virus at an MOI of 0.5 for 1 h. Infected cells were cultured for another 24 h with conventional medium, then treated with Trizol. The infection of authentic SARS-CoV-2 virus was quantified by real-time qPCR of RNA abundance.

### Statistical analysis

Statistical analysis of all data apart from CRISPRa screening was performed using GraphPad Prism software. The statistical significance was evaluated using Student’s *t* test and determined as *p* < 0.05. *P*-values were indicated in each of figure legends.

---

**fig. S1.**
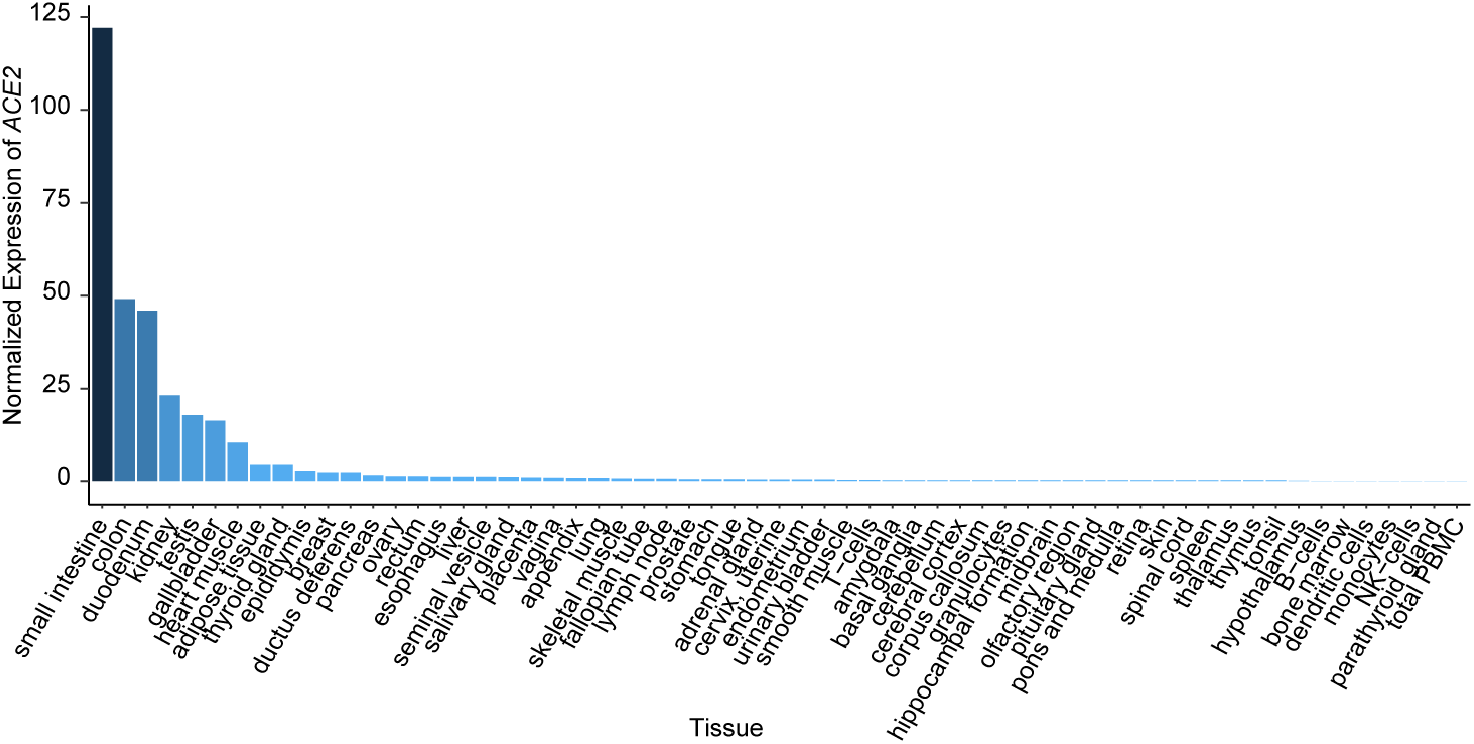
Normalized *ACE2* expression levels in different human tissues. The data used for analysis were retrieved from Human Protein Atlas normalized expression.

**fig. S2.**
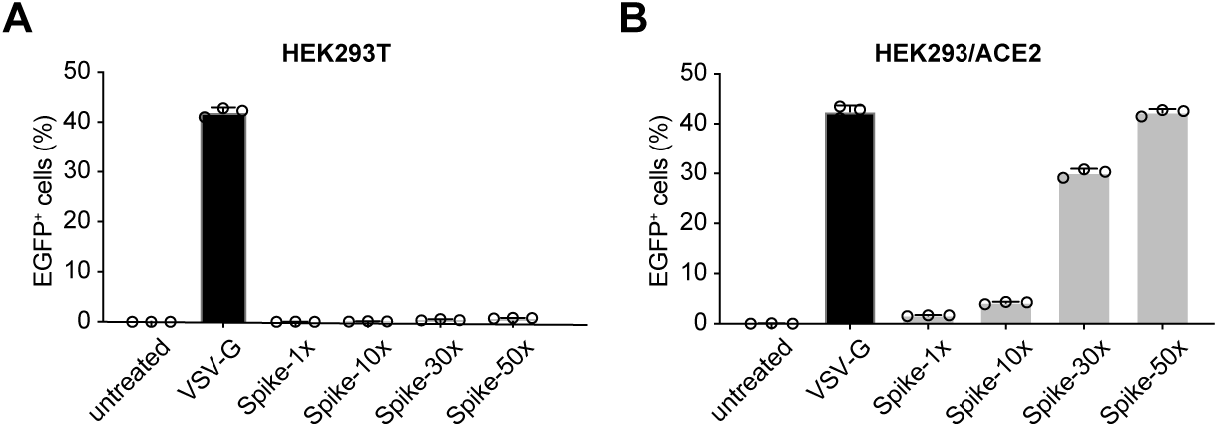
Examination of the approach in simulating SARS-CoV-2 infection using the SARS-CoV-2 pseudotyped virus. (**A**) Examination of the virus infection in wild type HEK293T cells using different concentrations of pseudovirus. HEK293T cells were respectively infected with 1-fold (Spike-1×), 10-fold (Spike-10×), 30-fold (Spike-30×) and 50-fold (Spike-50×) concentrated pseudovirus harboring SARS-CoV-2 S protein, and the EGFP^+^ percentages were analyzed by FACS 48 h post infection. The lentivirus harboring VSV-G protein was used for infection as a positive control following the same procedure. (**B**) Examination of the concentration of pseudovirus for achieving an efficient virus infection in HEK293 cells stably expressing *ACE2*.

**fig. S3.**
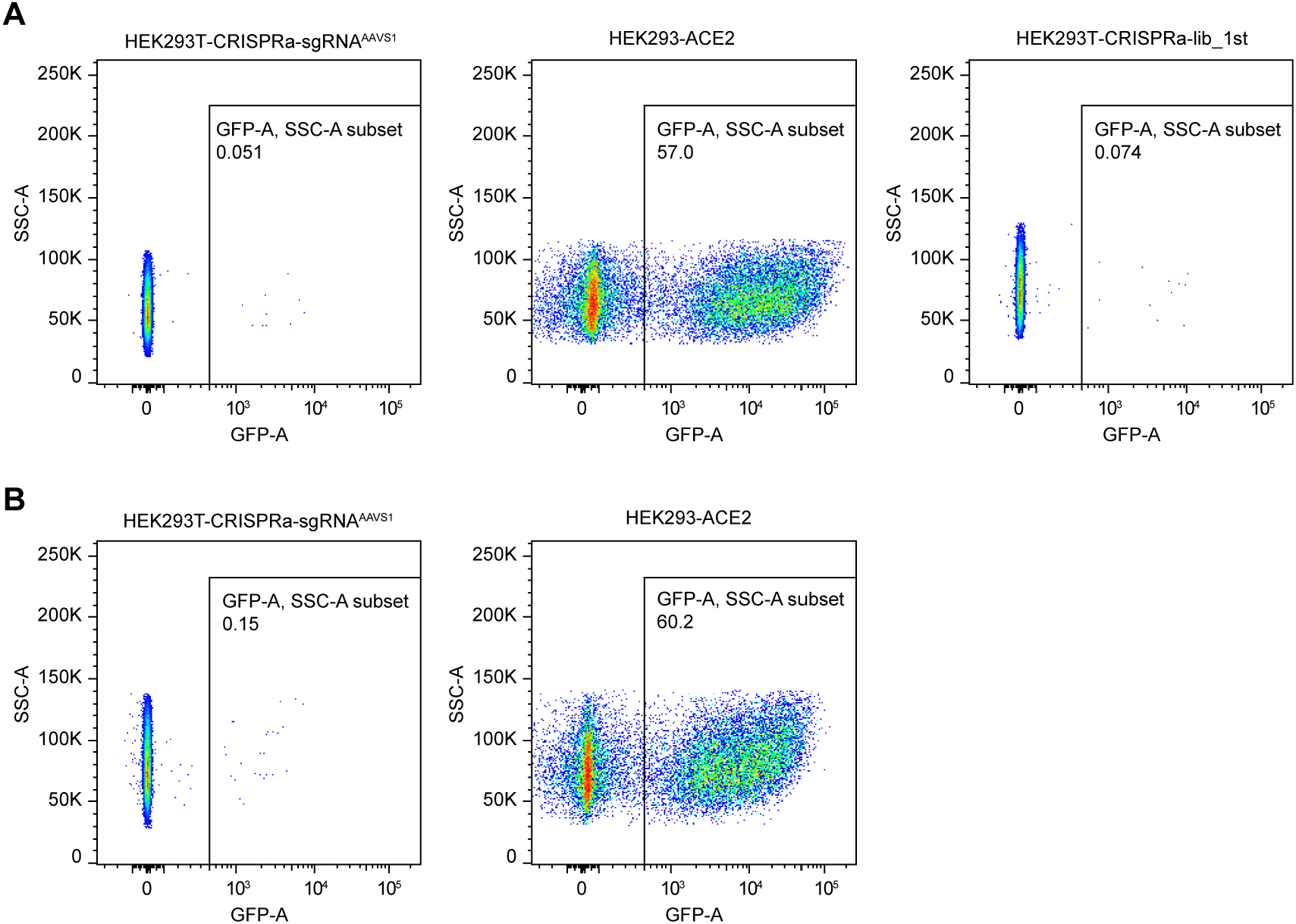
FACS selection of EGFP^+^ cells in each round of screening after SARS-CoV-2 pseudovirus infection. (**A**) The first FACS selection of EGFP^**+**^ cells from the HEK293T-CRISPRa library cells. Left: HEK293T-CRISPRa cells stably expressing *AAVS1*-targeting sgRNA were infected with SARS-CoV-2 pseudovirus (50-fold), serving as the negative control; Middle: HEK293 cells stably expressing ACE2 were infected with SARS-CoV-2 pseudovirus, serving as the positive control; Right: HEK293T-CRISPRa library cells were infected with SARS-CoV-2 pseudovirus for the first round. (**B**) The controls used in the second round of FACS selection of EGFP^**+**^ cells. The negative and positive controls were the same as in fig. S2A (left and middle), and the FACS selection of library cells for the second round was presented in Fig.1D.

**fig. S4.**
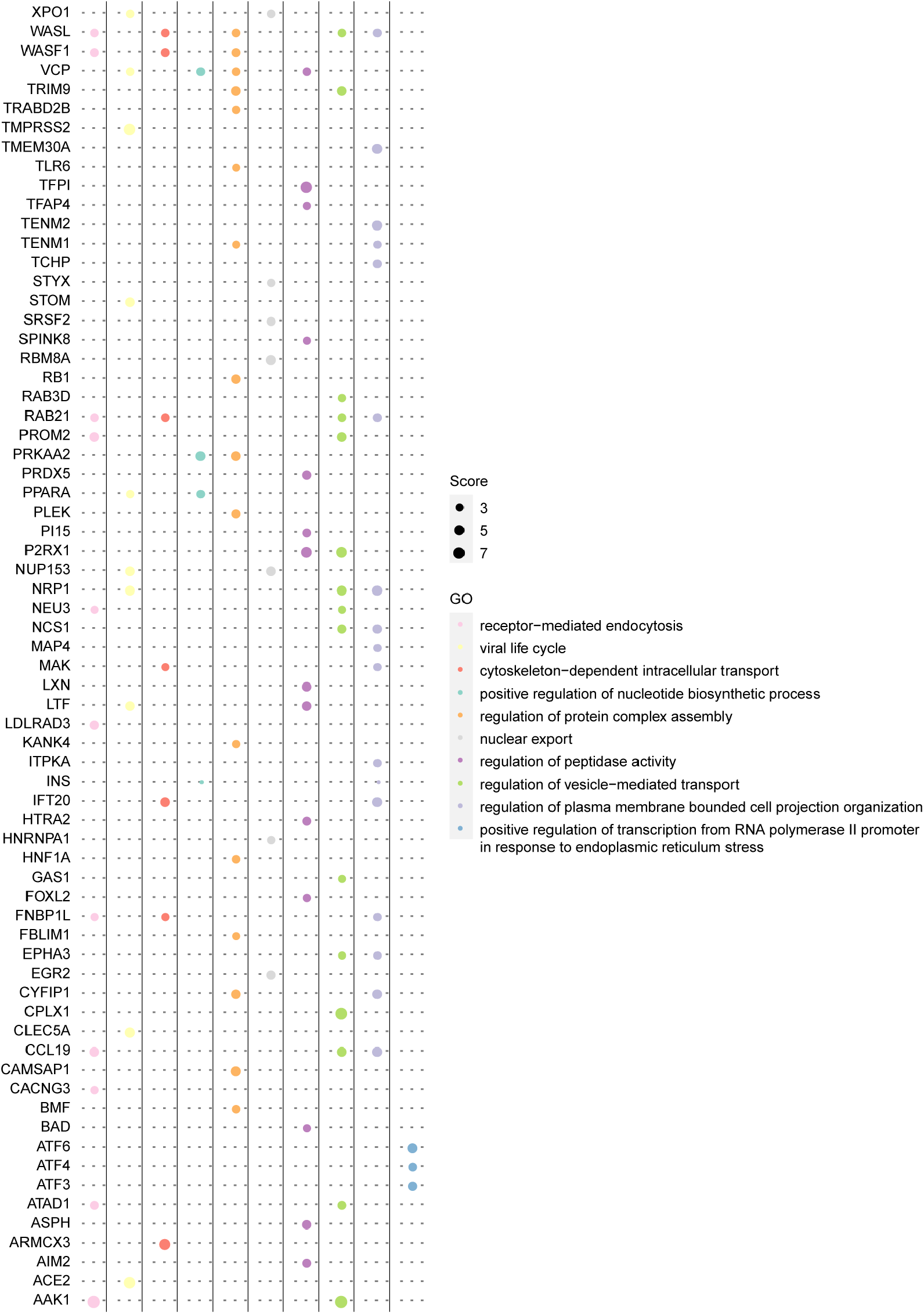
The total gene list from GO enrichment analysis. The size of round dots indicated scores of CRISPRa screening.

**fig. S5.**
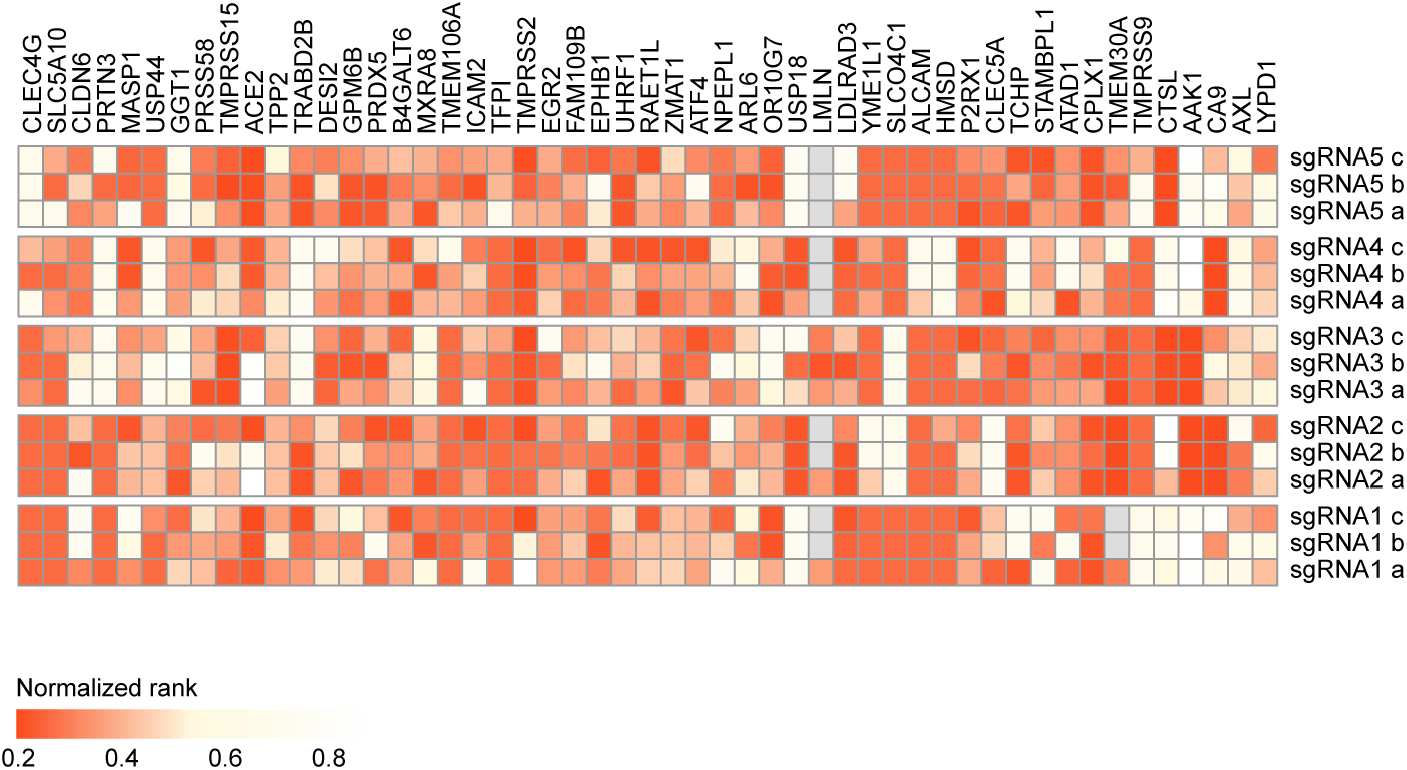
The performance of all the sgRNAs^eBAR^ targeting the identified candidates. Three eBARs of each sgRNA were indicated by a, b and c.

**fig. S6.**
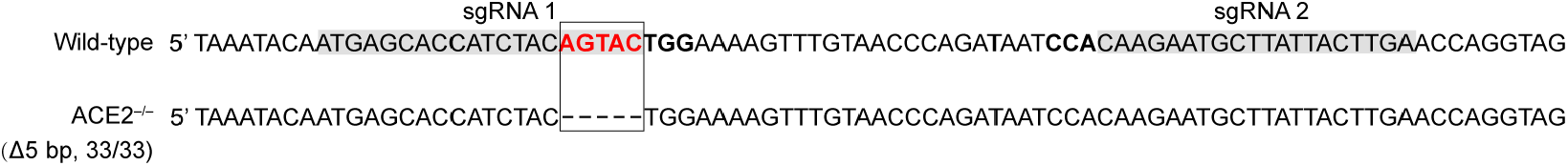
Partial coding sequences of the *ACE2* gene in the genome containing the sgRNA binding regions. The sequencing analysis of the mutated alleles were obtained from 33 randomly selected clones. The dashes indicate deletions.

**fig. S7.**
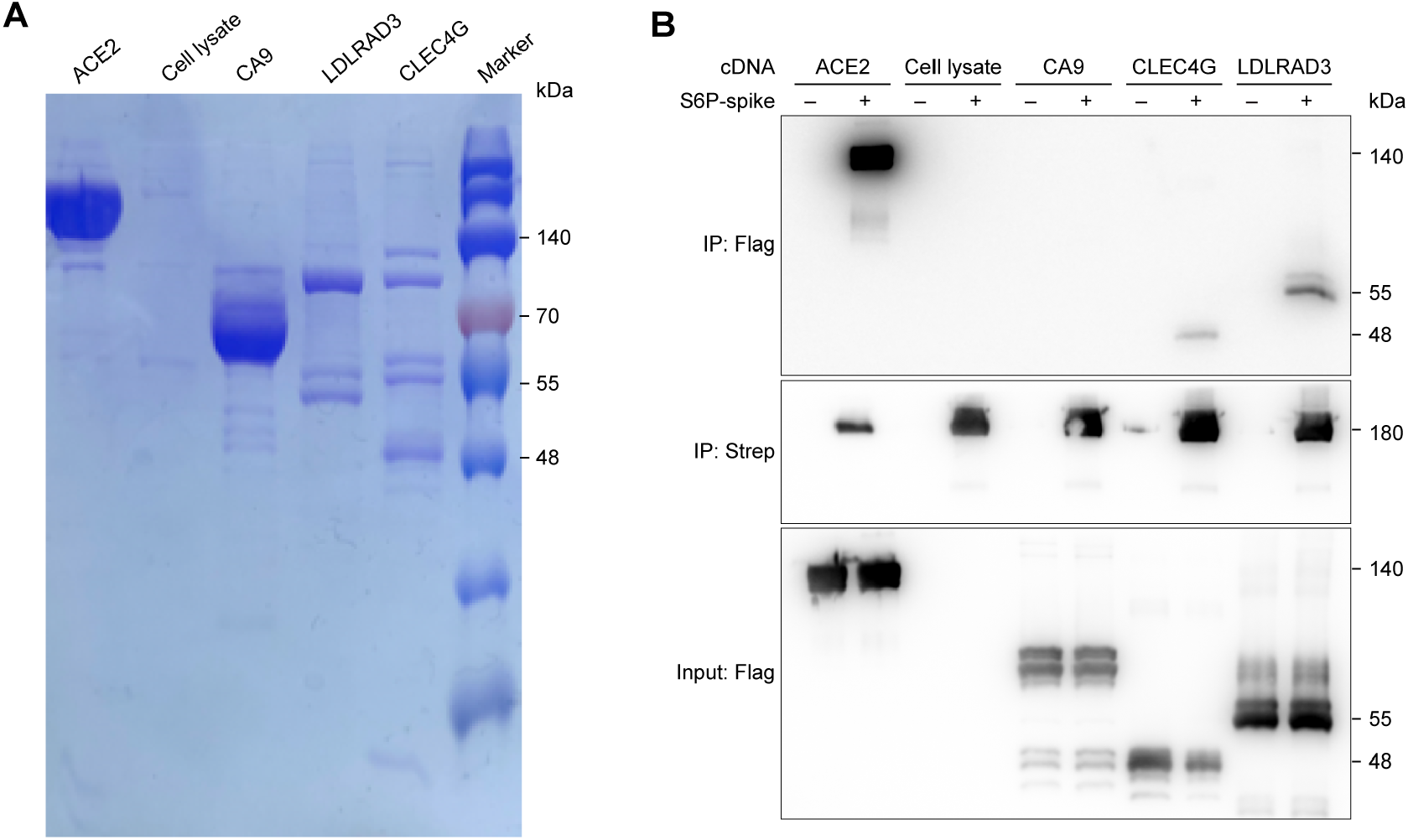
Direct binding of identified proteins to SARS-CoV-2 S. (**A**) Flag-tagged CA9, LDLRAD3, CLEC4G and ACE2 were purified and shown on a Coomassie blue-stained SDS-PAGE gel. (**B**) *In vitro* pull-down assay of purified ACE2, CA9, CLEC4G and LDLRAD3 to SARS-CoV-2 S. Strep-tagged SARS-CoV-2 S and FLAG-tagged candidate receptors were expressed in HEK293T cells and affinity-purified. Immunoblot analysis was conducted using anti-Flag and anti-Strep antibodies.

**fig. S8.**
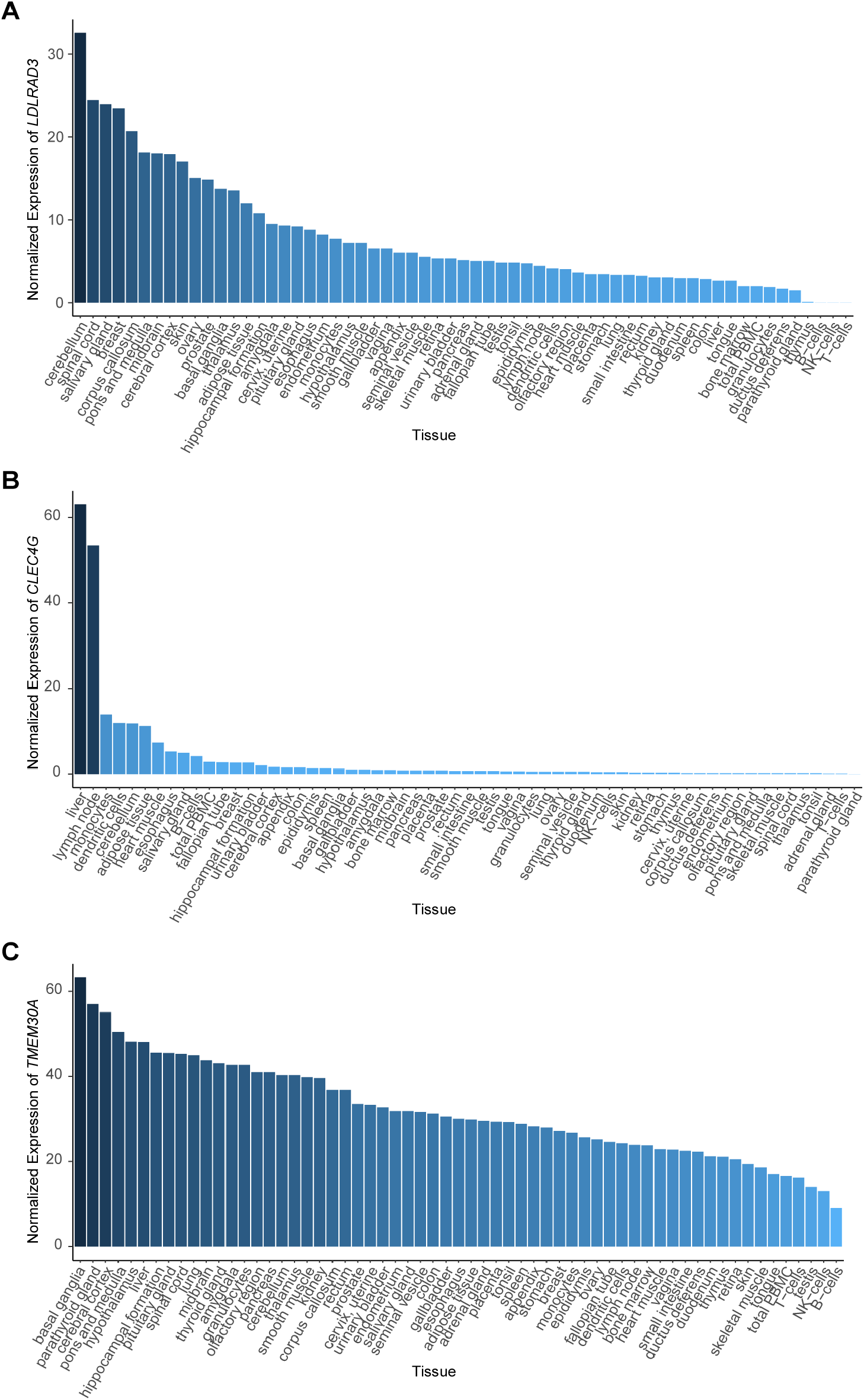
Expression patterns of candidate receptors within human tissues. The mRNA levels of *LDLRAD3* (**A**), *CLEC4G* (**B**) and *TMEM30A* (**C**) within human tissues were analysed using data retrieved from Human Protein Atlas.

## Acknowledgements

We acknowledge the National Center for Protein Sciences (Beijing) at Peking University for their assistance with fluorescence-activated cell sorting, particularly Dr. J.L., Ms H. Y., and Ms L.D. for their technical help. We acknowledge Dr. Ying Yu (Peking University) for her assistance in preparing the NGS library.

## Author Contributions

W.W. conceived and supervised this project. W.W., S.Z., Y.L., and Z.Z. (Zhou) designed the experiments. S.Z. and Y.L. performed the CRISPRa screening and the following validations with the help from A.C., F.T., Y.X., C.W., Q.L., X.N., and Q.P.. X.X. performed the authentic SARS-CoV-2 virus infection with the help from X.D., under the supervision of J.W.. Z.Z. (Zhang) performed the protein purification and pull-down assay with the help from S.C. and S.D., under the supervision of J.X.. Z.L. performed the bioinformatics analysis. S.Z., Y.L., Z.Z. (Zhou), and W.W. wrote the manuscript with the help of all other authors.

## Author Information

The authors declare no competing financial interests. Readers are welcome to comment on the online version of the paper. Correspondence and requests for materials should be addressed to W.W., J.W. and J.X.

